## Supplemental File for "IDH1 regulates human erythropoiesis by eliciting chromatin state reprogramming"

**Supplementary methods**

**Human Samples:** Human bone marrow samples were obtained from 10 AML/MDS patients with IDH1 mutation (4 MDS and 6 AML) in the Department of Hematology, First Affiliated Hospital of Zhengzhou University (Zhengzhou, China). Details of the samples are provided in Supplementary Table 1. Written informed consent was obtained from all participants, and the protocols were approved by the Ethics Committee of The First Affiliated Hospital of Zhengzhou University (2021-KY-0575-002).

**Generation of cell lines:** The K562 and HEL cell lines were obtained from The State Key Laboratory of Medical Genetics & School of Life Sciences, Central South University (Changsha, China), Human HUDEP2 cell line was obtained from Institute of Hematology & Blood Diseases Hospital, Chinese Academy of Medical Sciences & Peking Union Medical College (Tianjin, China). The detailed methods have been described previously including culture medium composition, the culture protocol.[1]

Nuclear IDH1 was deleted in the HUDEP2 cell line using the following approach: sgRNA sequences targeting different genomic regions of IDH1 (IDH1-sgRNA-Top: CACCGATGTAGATCCAATTCCACGT and IDH1-sgRNA-Bottom: AAACACGTGGAATTGGATCTACATC) were annealed and ligated with pL-CRISPR.EFS.GFP vector by restriction digest with BsmBI [2]. Production of lentiviral vectors was performed according to standard protocol. HUDEP2 cells were then infected with viral particles carrying sgRNA sequences targeting IDH1 and sorted with GFP to get Sg-IDH1 HUDEP2 cells.

pBabe-puro-NES-IDH1 was generated using primers encoding for the PKI NES (LALKLAGLDI) [3]. Forward (TCTAGGCGCCGGCCGGATCCCGCCACCATGTCC-AAAAAAATCAGTGGCGG) and reverse (ACCACTGTGCTGGCGAATTCTTAAA-TATCCAGGCCCGCCAGTTTCAGCGCCAGGTCGACAAGTTTGGCCTGAGCTA GTTT) primers were annealed and ligated with pBabe-puro by restriction digest with BamHI and EcoRI [4]. Production of lentiviral vectors was performed according to standard protocol. Sg-IDH1 HUDEP2 cells were infected with viral particles of pBabe-puro-NES-IDH1 and selected with Puromycin to get Sg-NES-IDH1 HUDEP2 cells.

**Protein extraction.** Total protein extraction: Cells were harvested and washed with PBS. Then, cells were centrifugated at 300×g for 5 minutes. Cell lysates were prepared with RIPA lysis buffer (#89900, Thermo Fisher Scientific) in the presence of proteinase inhibitor phenylmethanesulfonylfluoride fluoride (PMSF) (#36978, Thermo Fisher Scientific).

Cytoplasm and nuclear protein extraction were used by NE-PER™ Nuclear and Cytoplasmic Extraction Reagents kit (#78833) from Thermo Scientific™. The detail method was described as follows: Cells (5-10×$10^6$) were harvested and washed three times with PBS. Then fresh cells were centrifugated at 300×g for 5 minutes. Ice-cold CER I was added to the cells and the tube was vigorously vortexed on the highest setting for 15 seconds to fully suspend the cell pellet. Then cells were incubated on ice for 10 minutes. Ice-cold CER II was then added to the tube and vortexed for another 5 seconds on the highest setting. Then cells were incubated on ice for 2 minutes. The supernatant was collected after centrifugation for 5 minutes at 16,000×g and then immediately transferred to a clean pre-chilled tube. Cytoplasmic protein was fully extracted. Then, insoluble fractions were suspended using ice-cold NER and vortexed on the highest setting for 15 seconds. Samples were placed on ice and oscillated for 15 seconds every 10 minutes for 40 minutes. Finally, the supernatant containing extracted nucleoprotein was immediately transferred to a clean pre-chilled tube. All reagents used during the experiment should be placed on ice. Maintain the volume ratio of CER I: CER II: NER reagents at 200:11:100 μL, respectively. Protein concentration was measured by BCA protein concentration determination using Micro BCA™ Protein Assay Kit (23235, Thermo Fisher Scientific)

**Cytospin assay.** Obtaining sample slides: cells (5×$10^4$) were collected in a 1.5 mL Eppendorf tube (EP), washed with PBS, and centrifugated at 300×g for 5 minutes at 4 °C. Cells were resuspended with 200 μL PBS. Then, cells were adhered to slides by cell spin centrifugation. The fresh cytospin were fetched and dried carefully without damaging.

Proceeding with May Grunwald/Giemsa staining: Cells were stained with May-Grunwald solution (MG500, Sigma) for 5 minutes and then washed with 40 mM Tris buffer (pH 7.2) for 90 seconds. Then, they were dyed with Giemsa solution (GS500, Sigma) for 15 minutes at once. The images were finally taken using a standard light microscope (Axio Imager.A2, Carl Zeiss Microscopy GmbH, Jena, Germany).

**RNA extraction and quantitative reverse transcription-PCR assays.** Total RNA was separated from human erythroblasts at distinct stages using RNA extract kits (#74104, Tiangen Biotech). The concentration was established using the Nanodrop (Thermo Fisher Scientific). RNA samples underwent reverse transcription with HiFi-MMLV cDNA Kit (CW0744M, CWBIO). Quantitative reverse transcription-PCR was completed by using SYBR™ Select Master Mix (#4472903, Thermo Fisher Scientific) and Light Cycler 480 system (Roche Life Science). Primers were obtained from Harvard primer bank. Relative expression levels were normalized to GAPDH. The primer sequences of *IDH1* and *GAPDH* were shown in Supplemental Table 3.

**Flow cytometric analysis and Fluorescence-activated cell sorting of erythroblasts.** The differentiation of erythroid cells was assessed by the expression of surface markers using flow cytometer [5]. The erythroid cells at the distinct developmental stage were sorted using a MoFlo Astrios Cell Sorter (Beckman Coulter Life Science) as previously described [6].

**Immunofluorescence imaging microscopy.** Cells ($0.5\times10^6$) were collected and washed with 1×PBS. Then, cells were fixed with 1% paraformaldehyde for 15 minutes and permeabilized with 0.1% Triton X-100 in 0.25% paraformaldehyde-PBS for 10 minutes. Then, cells were incubated in PBS with 10% horse serum and 0.1% Triton X-100 for 30 minutes to minimize nonspecific antibody binding. Cells were incubated with primary antibodies at 4°C overnight, washed three times with PBS, and incubated with the appropriate secondary antibody at room temperature for 30 minutes. Nuclei were stained with Hoechst 33342 (blue). After washing three times with PBS to remove nonspecific staining. The cells were collected and seeded onto Thermo Scientific™ Nunc™ Lab-Tek™ II chamber (#155382). Images were collected and visualized under a confocal laser scanning microscope with a ×100 oil objective lens (Zeiss LSM780, Carl Zeiss Microscopy GmbH, Jena, Germany).

**Measurement of ROS.** DCFH-DA (2,7-Dichlorodihydrofluorescein diacetate) was used to assess reactive oxygen species (ROS) formation using Reactive Oxygen Species Assay Kit (S0033S, Beyotime Biotechnology). Cells ($0.5\times10^6$) were collected and washed with PBS buffer. Then, cells were resuspended and incubated in medium containing 10μM DCFH-DA at 37°C for 30 minutes. Cells were washed with PBS three times to remove DCFH-DA that had not entered cells. After entering cells, the DCCFH-DA probe was hydrolyzed by esterase to form DCFH, which was further oxidized by ROS in cells to form fluorescent DCF. The fluorescence intensity was measured using BD LSRFortessa™ flow cytometry to measure the level of ROS.

**Quantification of α-KG level.** α-KG were detected by using an α-KG assay kit (MAK054, Sigma), Preparation of standard substance: α-KG stock solution was diluted with water to 1mm standard buffer. 0, 2, 4, 6, 8, and 10 μL of the 1 mM α-KG standard buffer were added successively into a 96 well plate, generating 0 (blank), 2, 4, 6, 8, and 10 nmol/well standards. α-KG assay buffer was added to each well to bring the volume to 50μL. Sample Preparation: Cells ($0.5 \times 10^6$) were homogenized in 100μL ice cold α-KG buffer, then were centrifugated at 13,000×g for 10 minutes to remove insoluble material. Samples were prepared to a final volume of 100 μL reagent, including 94μL α-KG assay buffer, 2μL α-KG converting enzyme, 2μL α-KG development enzyme mix and 2μL flourescent peroxidase substrate. Blank control group was similar with the sample preparation group with no α-KG converting enzyme. Then the samples were incubated at 37°C for 30 minutes. SpectraMax i3 (Molecular Devices) was used to check the concentration of colorimetric product at 570 nm wavelength to determine the total α-KG concentration.

**Transmission electron microscopy.** Cells ($20\times10^6$) were collected and fixed with 2.5% glutaraldehyde. The fixed cells were washed in 0.1 M sodium caccodylate buffer (pH 7.2), and fixed with 1% osmium tetraoxide in sodium caccodylate buffer. Then, cells were dehydrated in ethanol (successively in 70%, 95%, and absolute ethanol), treated with propylene oxide (a transitional solvent), infiltrated in a mixture of propylene oxide and resin (Epon), embedded in pure resin mixture, and cured at 60°C. Thin sections of 50 nm were generated using an LKB 2088 Ultratome V, applied on copper slot grids and stained separately with 5% uranyl acetate and 3% lead citrate. Images were generated using a 120-Kv H-7700 Hitachi transmission electron microscope. The area of nuclear, heterochromatin and euchromatin were quantified using Image J software.

**RNA-seq and bioinformatics analysis.** Pre-sequencing experimental procedures:

Cells (2×10$^6$) were collected and washed twice by PBS. RNA was extracted and reverse transcribed into cDNA. The cDNA libraries were prepared using NovaSeq 5000/6000 S4 Reagent Kit (#A00358, Illumina) and sequenced on an Illumina X-ten (E00487, Illumina). The useful Perl script was used to filter the original data (Raw Data) and compare data to the reference genome by using HISAT2 which download from ENSEMBL database (http://www.ensembl.org/index.html, human genome builds GRCh38.87) [7]. Reads Count for each gene in each sample was counted by HTSeq v0.6.0 [8], and FPKM (Fragments Per Kilobase Million Mapped Reads) was then calculated to estimate the expression level. DESeq2 version 1.6.3 (https://bioconductor.org/packages/3.0/bioc/html/DESeq2.html) are package for Identifying Differentially Expressed Genes (DEGs) from data (FPKM > 10, q < 0.05, and fold change > 2) [9]. Gene Ontology (http://geneontology.org/) was used to make function enrichment analysis with $q < 0.05$. Gene Set Enrichment Analysis (GSEA) was performed using GSEA version 4.1.0 software (http://www.gsea-msigdb.org/gsea/downloads.jsp) [10]. RNA-seq data have been deposited in the GEO under accession code GSE223141.

**Assay for Transposase-Accessible Chromatin with high throughput sequencing.** Pre-sequencing experimental procedures: Cells (2×10$^6$) were collected and washed twice by PBS. Cells were prepared for detection, DNA transposition, PCR amplification, fragment selection, library quality control, and computer sequencing. Sequencing and analysis: Filter data to get high-quality existing data. After the quality control test is qualified, clean data will be mapped the reference genome which download from ENSEMBL database (http://www.ensembl.org/index.html, human genome builds GRCh38.87) using Bowtie2 (https://sourceforge.net/projects/bowtie-bio/files/bowtie2/2.3.5.1/) [11]. Identifying statistically significant differentially accessible regions using DEseq2 parameters of DiffBind version 3.8.4 (https://bioconductor.org/packages/release/bioc/html/DiffBind.html) [12]. The ATAC peak/region <1000bp from the nearest TSS was designated as the promoter. Reads coverage and depth were calculated by samtools version 1.6 software (https://github.com/samtools/samtools/releases/) [13]. Signal track files in Big Wig format were generated using the Deeptools version 2.5.7

(https://deeptools.readthedocs.io/en/develop/content/installation.html) and were normalized to 1 million reads for visualization [14]. MACS2 version 2.1.1 (https://pypi.org/project/MACS2/) was used to identify peaks using parameters "nomodel shift -100 extsize 200" with a q value of < 0.05 [15]. Annotation of peaks were performed using ChIPseeker version 1.20.0 software (https://www.bioconductor.org/packages/3.9/bioc/html/ChIPseeker.html) [16]. GO enrichment analysis and visualization of differentially accessible regions was performed using the clusterProfiler version 4.6.0 (https://bioconductor.org/packages/release/bioc/html/clusterProfiler.html) with q < 0.05 [17]. Overrepresented motif analysis was performed by Hypergeometric Optimization of Motif Enrichment (HOMER) tool Homer version 4.11 (http://homer.ucsd.edu/homer/) [18]. ATAC-seq have been deposited in the GEO under accession code GSE222401.

**Chromatin immunoprecipitation and sequencing.** Sample preparation procedures: cells ($20 \times 10^6$) were collected and washed twice by PBS, then cells were resuspended in 1% formaldehyde, cross-linked and quenched with 125mm glycine in a shaker at 25°C for 10 minutes. Cells were centrifugated 1500×g at 25°C for 5 minutes. The precipitates were suspended using pre-cooled PBS and centrifuged at 1500×g for 5 minutes. Supernatants which around the sediment were removed very carefully.

Pre-sequencing experimental procedures: Firstly, nanodrop, qubit and Agilent 2100 were used to assess the sample quality, including purity, concentration and fragment distribution. For the library building process, DNA fragments were repaired at the end, added with basic group A and sequencing connector; then PCR amplification and fragment screening were used to complete the preparation of the library. The qualified library was ready for sequencing.

Sequencing and analysis: ChIP-seq reads were aligned to the reference genome using Bowtie2 version 2.3.5.1 (https://sourceforge.net/projects/bowtie-bio/files/bowtie2/2.3.5.1/), and only uniquely and non-duplicate mapped reads were utilized to perform the downstream analysis. MACS2 version 2.1.1 (https://pypi.org/project/MACS2/) was used to identify peaks using parameters "–broad–broad-cutoff 0.01" with a q value of < 0.05. Annotation of peaks were performed using ChIPseeker version 1.20.0 software (https://www.bioconductor.org/packages/3.9/bioc/html/ChIPseeker.html). KLF1 raw ChIP-seq data were downloaded from the Gene Expression Omnibus (GEO) GSE104574[19]. Other analysis methods as described for ATAC-seq. ChIP-seq data have been deposited in the GEO under accession code GSE222296.

**Supplementary Figures**

**Supplemental Figure 1.The expression level of IDH1 during erythropoiesis. (A)** RNA-seq analysis of IDH1 on each erythroid cells. **(B)** qRT-PCR results showing IDH1 expression in erythroblasts infected with lentivirus containing Luciferase-shRNA, IDH1-shRNA1, and IDH1-shRNA2 on D7, D11, and D15. **(C)** Representative images of Western Blotting showing IDH1 expression level in erythroblasts infected with Luciferase-shRNA, IDH1-shRNA1, and IDH1-shRNA2 on D7, D11, and D15.The results were normalized to GAPDH protein expression level. **(D)** Quantitative analysis of the knockdown efficiency of IDH1 from three independent experiments.

**Supplemental Figure 2. Deficiency of IDH1 slightly affect the proliferation and have on effect on apoptosis on terminal erythroblast. (A)** qRT-PCR results showing IDH1 expression in erythroblasts infected with lentivirus containing Luciferase-shRNA, IDH1-shRNA1, and IDH1-shRNA2 on days 7, 11, and 15. **(B)** Representative images of western blotting showing IDH1 expression level in erythroblasts infected with Luciferase-shRNA, IDH1-shRNA1, and IDH1-shRNA2 on day 7, day 11, and day 15. Results were normalized to GAPDH protein expression level. **(C)** Quantitative analysis of the knockdown efficiency of IDH1 from three independent experiments. **(D)** Growth curves of cells including Luciferase-shRNA, IDH1-shRNA1, and IDH1-shRNA2 at each day. **(E)** Representative flow cytometric profiles of apoptosis stained with 7AAD and Annexin V at day 13 and day 15 of culture cells transfected with Luciferase-shRNA, IDH1-shRNA1, and IDH1-shRNA2. **(F)** Quantitative analysis of apoptosis cells from three independent experiments. Statistical analysis is from 3 independent experiments, and the bar plot represents mean ± SD of triplicate samples. Not significant (ns), * $p < 0.05$, ** $p < 0.01$, *** $p < 0.001$.

**Supplemental Figure 3. siRNA mediated knockdown of IDH1 have no effect on apoptosis and GPA expression.** (A) Representative images of Western Blotting showing IDH1 expression level in erythroblasts with siRNA-NC, IDH1-siRNA1, and IDH1-siRNA2 on days 7,11, and 15. The results were normalized to GAPDH protein expression level. (B) Quantitative analysis of the knockdown efficiency of IDH1 from three independent experiments. (C) Quantitative analysis of apoptosis cells from three independent experiments. (D) Representative flow cytometry profiles of apoptosis stained with 7AAD and Annexin V on erythroid cells with siRNA-NC, IDH1-siRNA1, and IDH1-siRNA2 on day 7, day 11, and day 15. (E) Flow cytometric analysis showing the percentage of GPA-positive cells on day 7, day 11, and day 15. (F) Quantitative analysis of GPA-positive cells from three independent experiments.

**Supplemental Figure 4. Deficiency of IDH1 affect the generation of orthochromatic erythroblasts. (A)** Representative flow cytometry profiles of GPA staining of erythroid cells cultured on days 7, 9, 11, 13, and 15. **(B)** Quantitative analysis of the percentage of GPA-positive cells from three independent experiments. **(C)** Representative flow cytometry profiles of double-stained with band 3 and α4-integrin on days 7, 9, 11, 13, and 15. **(D)** Representative cytospin images on days 7, 9, 11, 13, and 15. Scale bar, 10 μm. **(E)** Quantitative analysis of the percentage of each stage cells from three independent experiments. Statistical analysis is from 3 independent experiments, and the bar plot represents mean ± SD of triplicate samples. Not significant (ns), * $p < 0.05$, ** $p < 0.01$, *** $p < 0.001$.

**Supplemental Figure 5. siRNA mediated knockdown of IDH1 impaired the terminal stage erythropoiesis. (A)** Flow cytometric analysis showing the expression of α4 integrin and band 3 of erythroid cells cultured for day 7, day 11, and day 15. **(B)** Quantitative analysis of the percentage of each stage cells from three independent experiments. (1) day 7, (2) day 11, (3) day 15. **(C)** Representative cytospin images on day 15 cultured erythroid cells. Scale bar, 10 μm. **(D)** Statistical analysis of the ratio of abnormal nuclear cells from three independent experiments. **(E)** Flow cytometric analysis showing the enucleation efficiency of Luciferase-shRNA, IDH1-siRNA1 and IDH1-siRNA2 on day 13 and day 15. **(F)** Statistics analysis of enucleation efficiency from three independent experiments. Statistical analysis is from 3 independent experiments, and the bar plot represents mean ± SD of triplicate samples. Not significant (ns), * $p < 0.05$, ** $p < 0.01$, *** $p < 0.001$.

**Supplemental Figure 6. Deficiency of IDH1 impaired nuclear condensation. (A)** Representative confocal images of proerythroblasts, basophilic erythroblasts, polychromatic erythroblasts, and orthochromatic erythroblasts. The nuclei was stained with DAPI and the color was blue. The cell membrane was stained with GPA and the color was red. Scale bar, 10 μm. **(B)** Quantitative analysis of the ratio of the nuclear and cytoplasm from three independent experiments.

**Supplemental Figure 7. IDH1 deficiency induced increase of ROS and decrease of α-KG. (A)** Flow cytometry analysis showed the level of ROS on day 15 erythroid cells with Luciferase-shRNA, IDH1-shRNA1, and IDH1-shRNA2. Quantitative analysis of ROS from three independent experiments. **(B)**Flow cytometry analysis showed the level of ROS on day 15 erythroid cells with Luciferase-shRNA, IDH1-shRNA1, and IDH1-GSH-50μM. Quantitative analysis of ROS from three independent experiments. **(C)**Flow cytometry analysis showed the level of ROS on day 15 erythroid cells with Luciferase-shRNA, IDH1-shRNA1, and IDH1-NAC-10 μM. Quantitative analysis of ROS from three independent experiments. **(D)**Quantitative analysis of the concentration of α-KG with Luciferase-shRNA, IDH1-shRNA1, and IDH1-shRNA2 on day 15 from three independent experiments. **(E)** Quantitative analysis of the concentration of α-KG after supplement α-KG (50 μM) on day 15 from three independent experiments. Statistical analysis is from 3 independent experiments, and the bar plot represents mean ± SD of triplicate samples. Not significant (ns), * $p < 0.05$, ** $p < 0.01$, *** $p < 0.001$.

**Supplemental Figure 8. IDH1 localizes to nucleus during human erythropoiesis. (A-B)** Representative images on location of IDH1 at erythroid differentiation of hemin-induced **(A)** K562 and **(B)** HEL cell lines. IDH1 (green), GPA (red) and Hoechst 33342 (blue). Scale bars, 5μm. IDH1 MFI of nucleus and cytoplasm of erythroid cells was shown at right panel. Data are presented as the mean ± SD from three independent experiments containing at least 30 cells each. **(C)** Representative immunofluorescence images of IDH1 (green), GPA (red) and Hoechst 33342 (blue) staining of the paraffin-embedded human bone marrow cells. a. AML-2, b. MDS-RAEB2, c. AML-3, d. MDS, e. AML-4, f. MDS-EB2, e. AML-5, h. AML-6. Scale bars, 5μm. IDH1 MFI of nucleus and cytoplasm of erythroid cells was shown at right panel. Data are presented as the mean ± SD from three independent experiments containing at least 30 cells each.

Supplemental Figure 9. Knockout nuclear IDH1 lead to cell number decrease of HUDEP2 cells. (A) Representative growth curve of control, sg-IDH1, sg-NES-IDH1， sg-PLVX-IDH1 on days 0, 2, 4, 6, 8. (B) Quantitative analysis of the concentration of α-KG from three independent experiments. (C) Representative flow cytometry profiles of apoptosis with Annexin V and 7AAD on days 8. (D) Quantitative analysis of the apoptosis percentage from three independent experiments. (E) Representative profiles flow cytometry of cell cycle with Edu and 7AAD on days 8. (F) Quantitative analysis of percentage of G0/G1, S and G2/M on days 8. Statistical analysis is from 3 independent experiments, and the bar plot represents mean  SD of triplicate samples. Not significant (ns), * $p < 0.05$, ** $p < 0.01$, *** $p < 0.001$.

**Supplemental Figure 10. IDH1 deficiency induced aberrant distribution and accumulation of histone modifications.**

**(A)** Representative Western Blotting images showed the total protein expression level of histone modification on day 15. **(B)** Quantitative analysis the relative protein expression level of histone modification on day 15.

**Supplemental Figure 11. The location of IDH1 and H3K79me3 in terminal erythroblasts of sg-NES-IDH1 HUDEP-2 cell line and AML/MDA patients. (A)**Representative immunofluorescence images of location of H3K79me3 in control and sg-NES-IDH1 HUDEP-2 cell lines. H3K79me3 (purple), GPA (red) and Hoechst 33342 (blue). Scale bars, 5μm. **(B)** Representative immunofluorescence images of IDH1 (green), H3K79me3 (purple), GPA (red) and Hoechst 33342 (blue) staining of the paraffin-embedded human bone marrow cells.

**Supplemental Figure 12. ATAC-seq analysis. (A)** Principal component analysis of (PCA) of ATAC-seq data. **(B)** Pearson correlation analysis of ATAC-seq data. **(C)** Bar plot displayed the distribution of peaks relative to gene features for IDH1-shRNA and Luciferase-shRNA.

**Supplemental Figure 13. RNA-seq analysis. (A)** Principal component analysis (left) and Pearson correlation analysis (right) of RNA-seq data. **(B)** The volcano map showed DEGs that upregulated (red color) and downregulated (blue color). **(C)** Representative images displayed DEGs of Luciferase-shRNA and IDH1-shRNA. **(D)** GO analysis. **(E)**

Upregulated genes in chromatin associated pathways in IDH1-shRNA group. The color represents log transformed adjusted p-value; the width indicates the number of DEGs in the category. **(F)** Heatmaps displayed DEGs associated chromatin of Luciferase-shRNA and IDH1-shRNA.

**Supplemental Figure 14. Integrated analysis of ChIP-seq, ATAC-seq and RNA-seq. (A)** Box plot showed that the genes modified by H3K79me3 were upregulated. **(B)** Heatmaps displayed DEGs with promoter region marked by H3K79me3. **(C)** The mean of ATAC signals at H3K79me3 modification sites.

**Supplemental Figure 15. Treatment with SIRT1 activator have no effet on cell differentiation and apoptosis of terminal erythroblasts.(A)**Representative flow cytometry profiles of GPA staining of with Normal-SRT1720 (0 nM, 100 nM, 500 nM, 2.5 μM) on days 9, 11, 13, and 15. **(B)** Quantitative analysis of the percentage of GPA-positive cells from three independent experiments. **(B) (C)** Representative flow cytometric profiles of apoptosis stained with 7AAD and Annexin V on day 15 erythroid cells with with Normal-SRT1720 (0 nM, 100 nM, 500 nM, 2.5 μM). **(D)** Quantitative analysis of apoptosis cells from three independent experiments. **(E)** Representative flow cytometric profiles of apoptosis stained with 7AAD and Annexin V at day 15 of culture cells with Normal-SRT1720 (0 nM, 100 nM, 500 nM, 2.5 μM). **(F)** Quantitative analysis of apoptosis cells from three independent experiments. Statistical analysis is from 3 independent experiments, and the bar plot represents mean ± SD of triplicate samples. Not significant (ns), * $p < 0.05$, ** $p < 0.01$, *** $p < 0.001$.

**Supplemental Figure 16. Treatment with SIRT inhibitor have no effet on cell differentiation and apoptosis of terminal erythroblasts.**

**(A)** Representative flow cytometry profiles of GPA staining of Luciferase-shRNA, IDH1-shRNA, IDH1-shRNA-EX527 (10 nM, 200 nM) cells on days 9, 11, 13, and 15. **(B)** Quantitative analysis of the percentage of GPA-positive cells from three independent experiments. **(C)** Representative flow cytometry profiles of double-stained with band 3 and α4-integrin on days 9, 11, 13, and 15. **(D)** Quantitativeanalysis of the percentage of each stage cells from three independent experiments. **(E)** Representative flow cytometric profiles of apoptosis stained with 7AAD and Annexin V at day 15 of culture cells with Luciferase-shRNA, IDH1-shRNA, IDH1-shRNA-EX527 (10 nM, 200 nM) cells. **(F)** Quantitative analysis of apoptosis cells from three independent experiments. Statistical analysis is from 3 independent experiments, and the bar plot represents mean ± SD of triplicate samples. Not significant (ns), * $p < 0.05$, ** $p < 0.01$, *** $p < 0.001$.

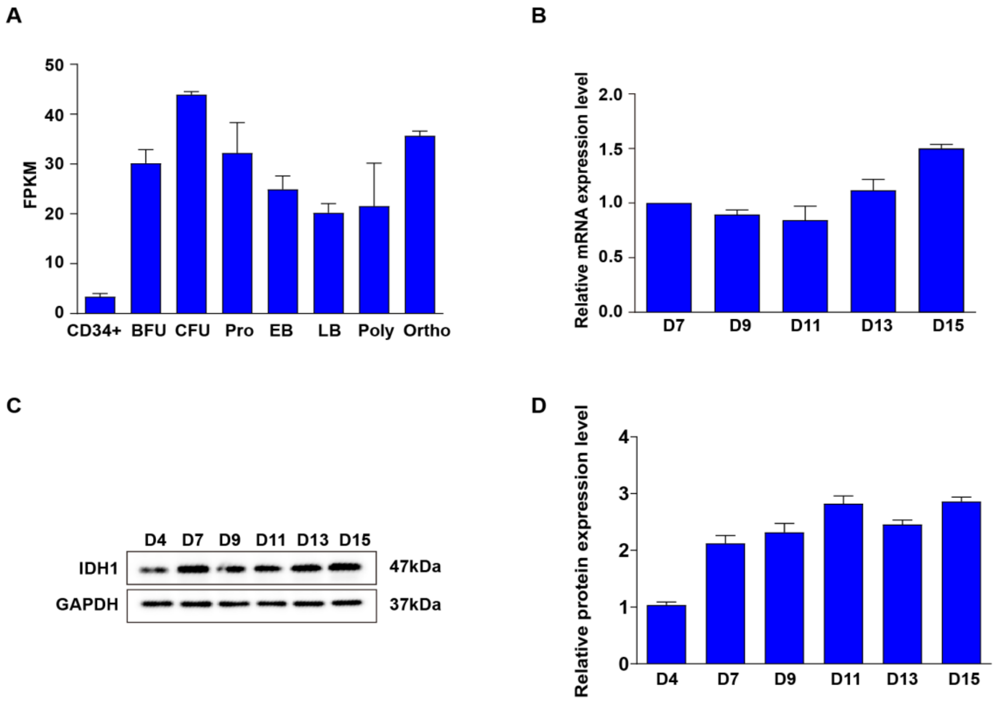


**Supplemental Figure 1.**The expression level of IDH1 during erythropoiesis.

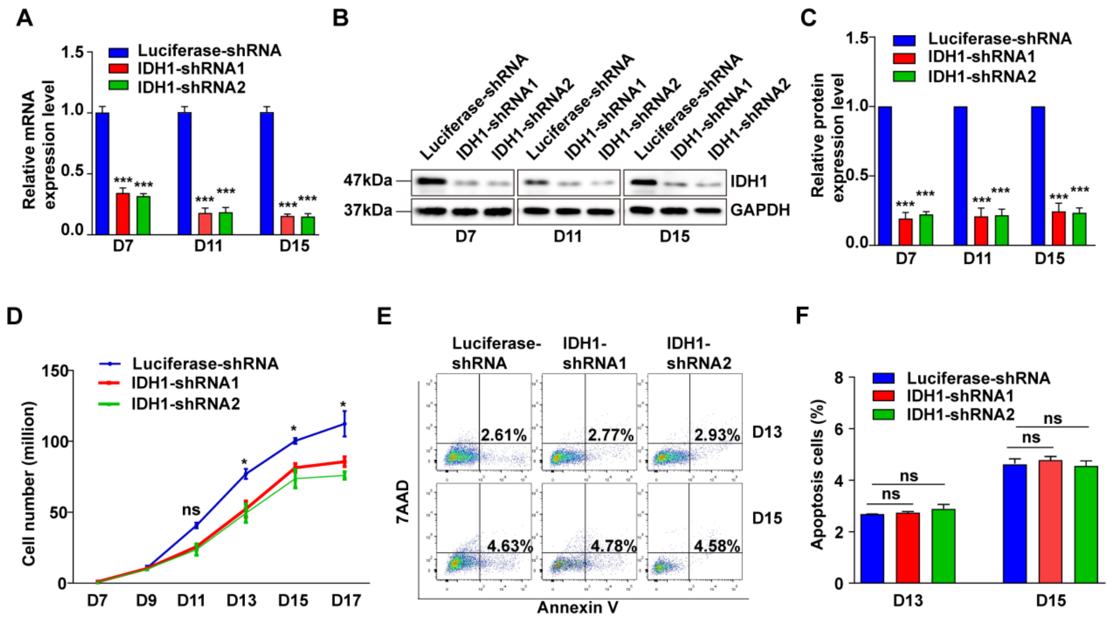

**Supplemental Figure 2.** Deficiency of IDH1 slightly affect the proliferation and have on effect on apoptosis on terminal erythroblast.

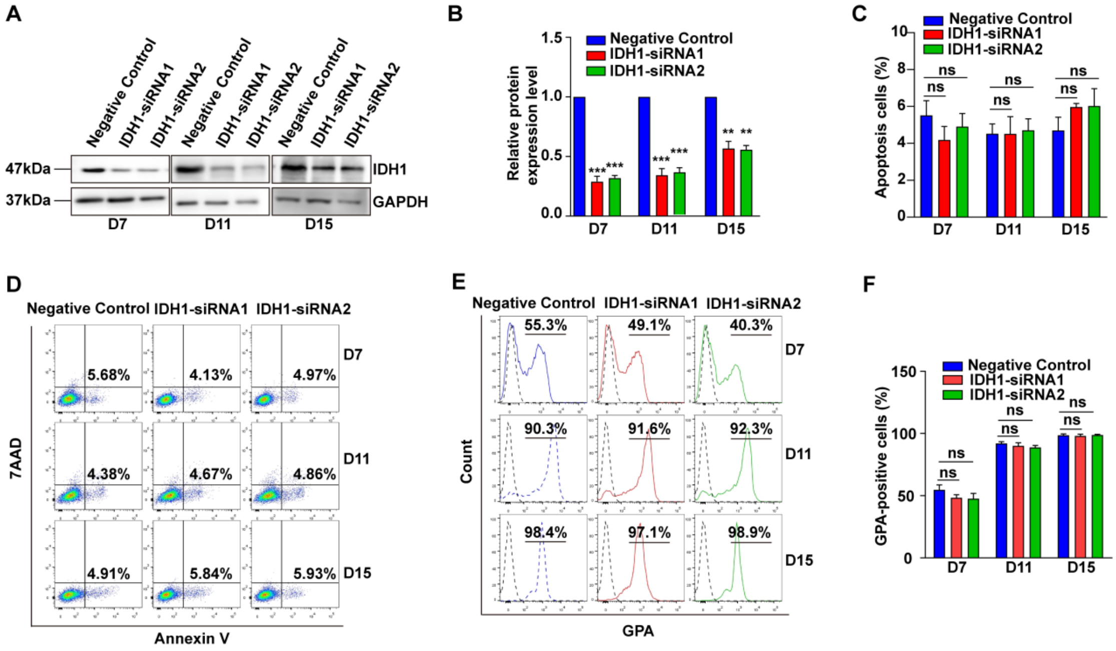

**Supplemental Figure 3.** siRNA mediated knockdown of IDH1 have no effect on cell growth and GPA expression.

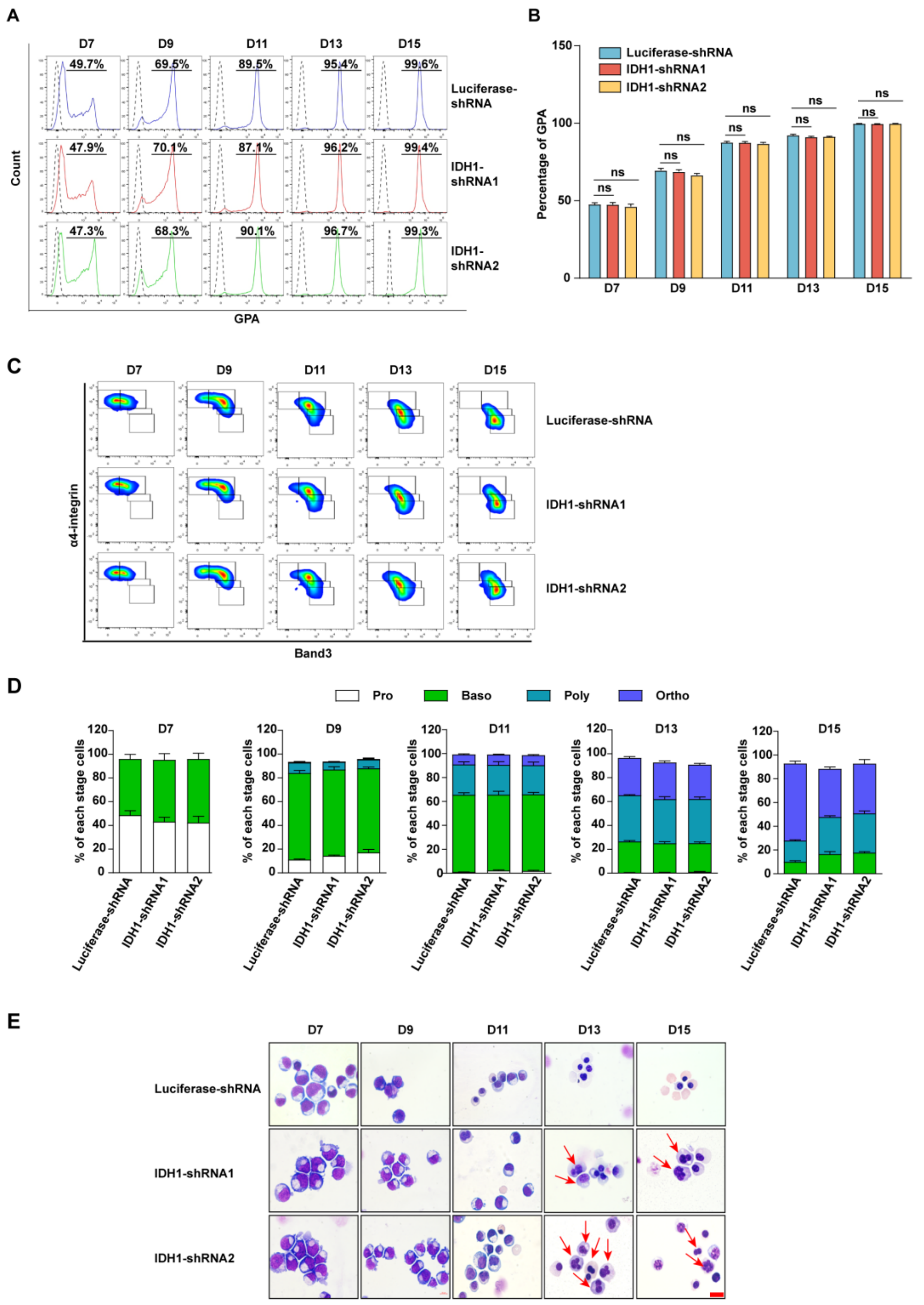


**Supplemental Figure 4.** Deficiency of IDH1 affect the generation of orthochromatic erythroblasts.

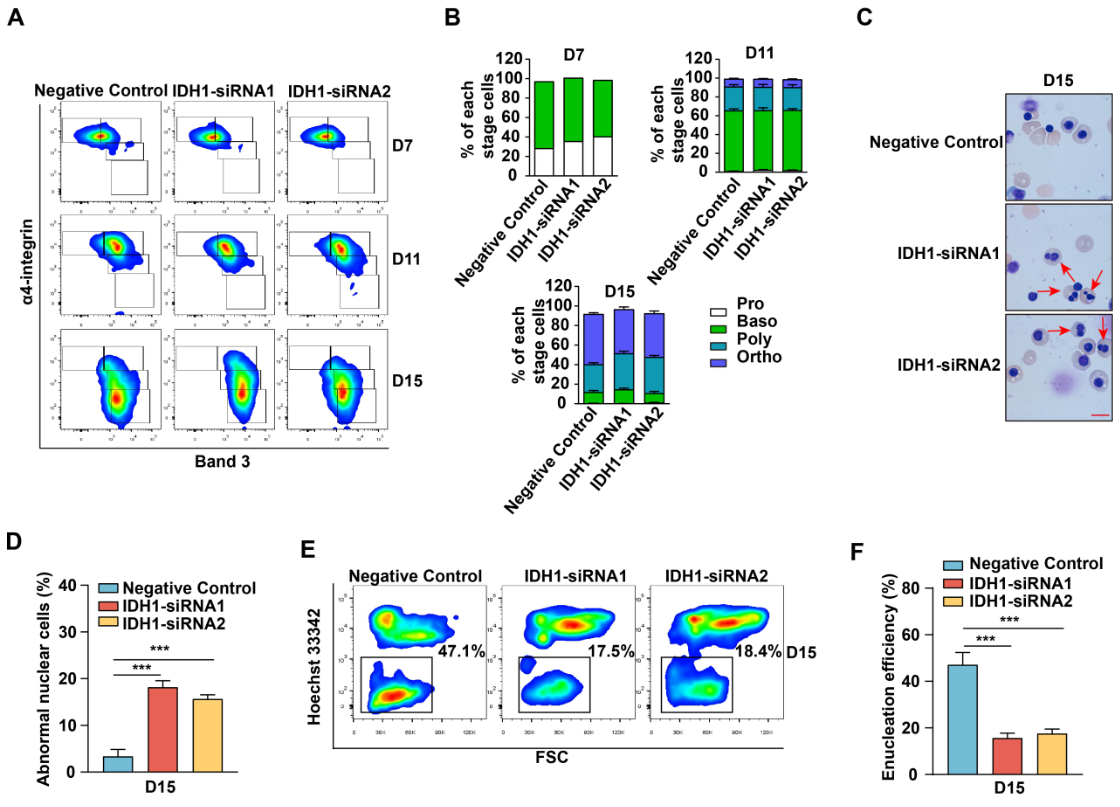


**Supplemental Figure 5.** siRNA mediated knockdown of IDH1 impaired the terminal stage erythropoiesis.

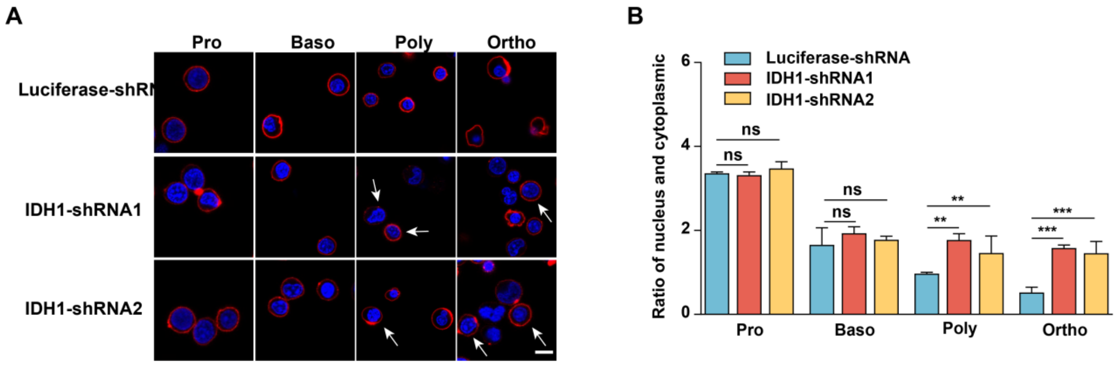


**Supplemental Figure 6.** Deficiency of IDH1 impaired nuclear condensation.

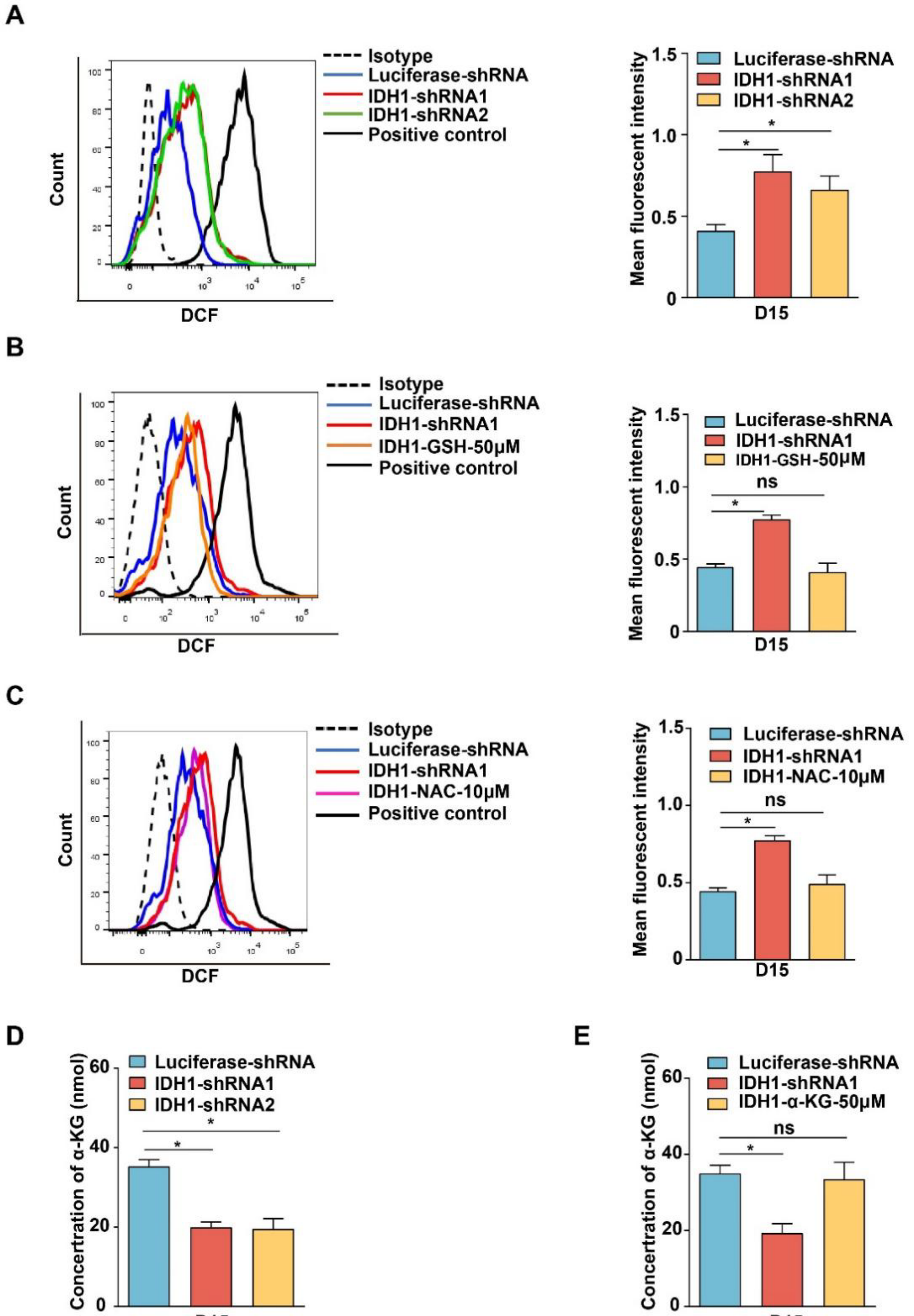


**Supplemental Figure 7.** IDH1 deficiency induced increase of ROS and decrease of α-KG

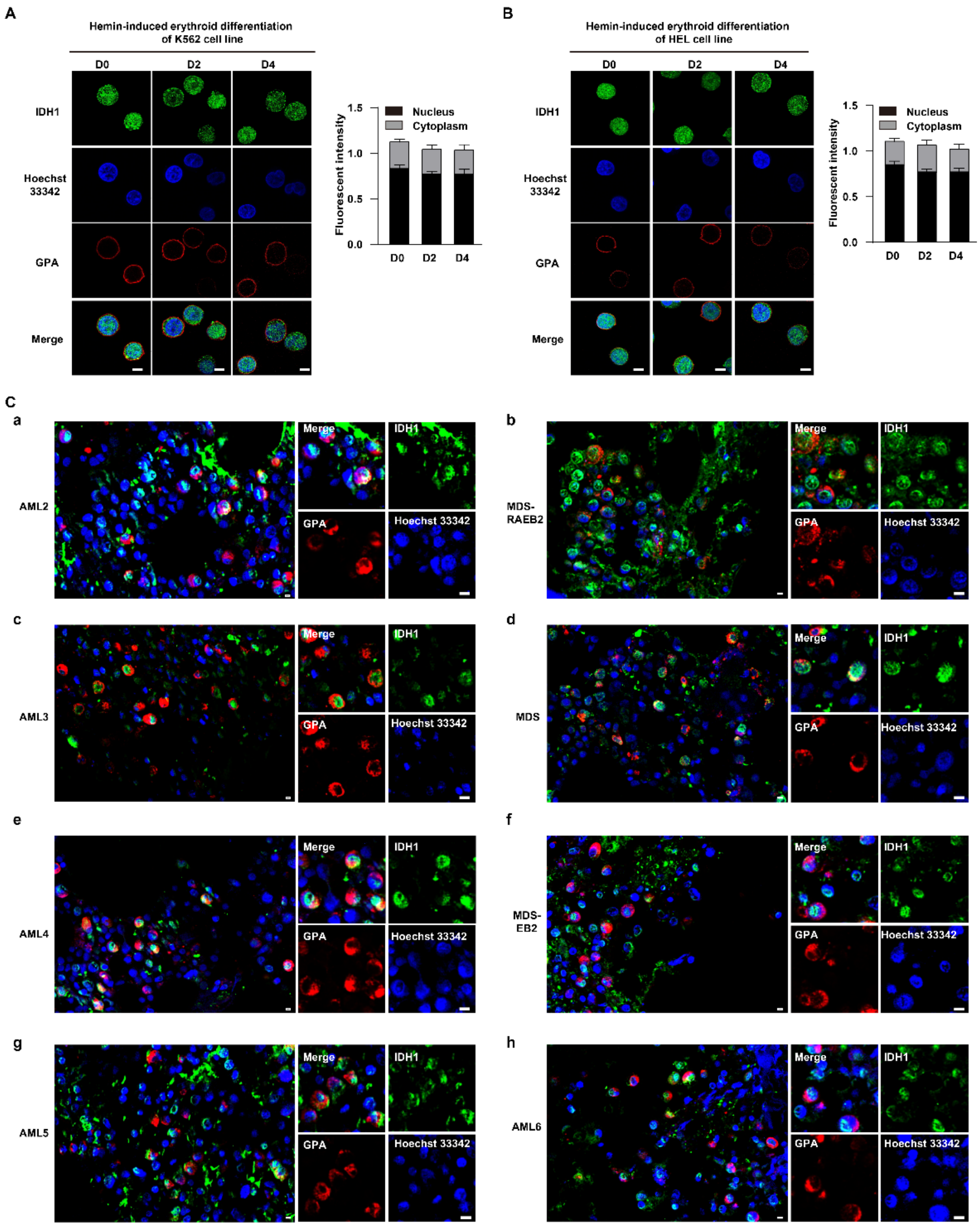


**Supplemental Figure 8.** IDH1 localizes to nucleus during human erythropoiesis.

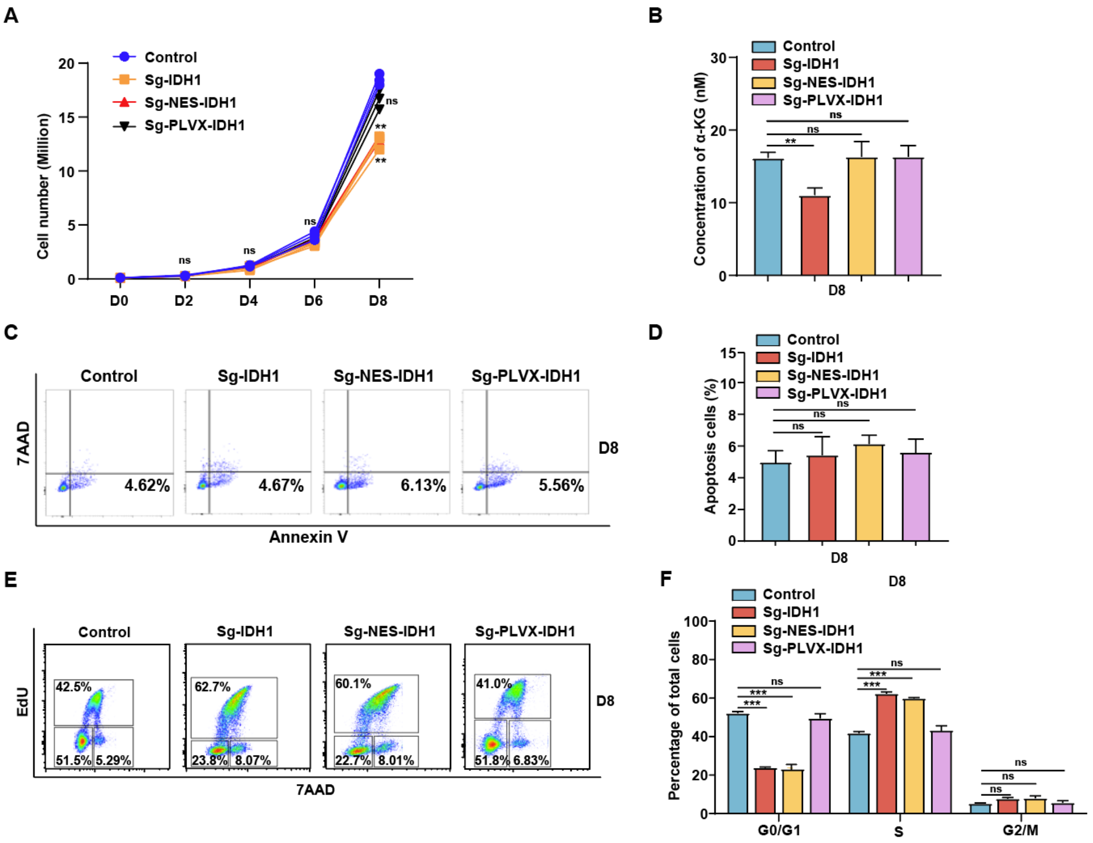


**Supplemental Figure 9.** Knockout nuclear IDH1 lead to cell number decrease of HUDEP2 cells.

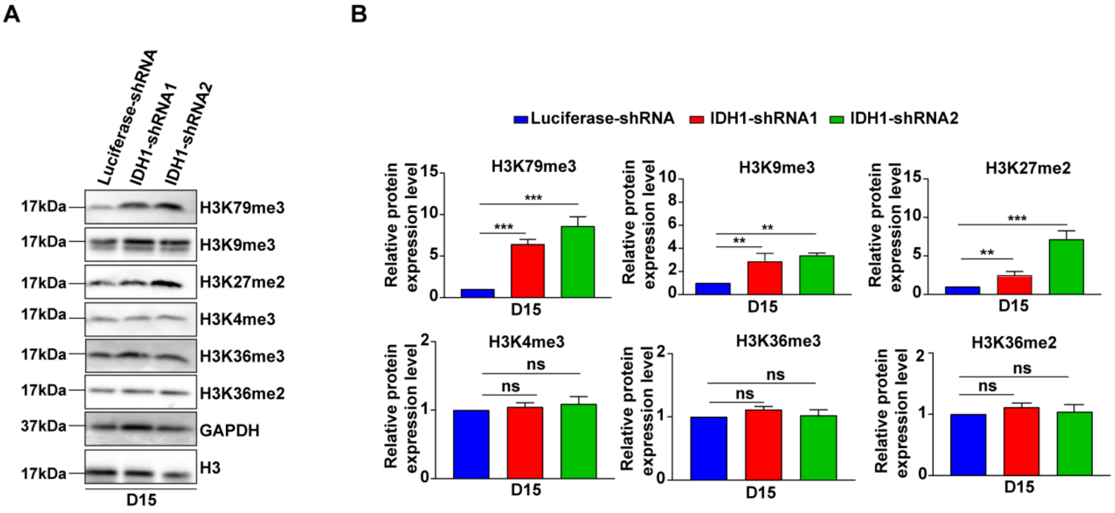


**Supplemental Figure 10.** IDH1 deficiency induced aberrant distribution and accumulation of histone modifications.

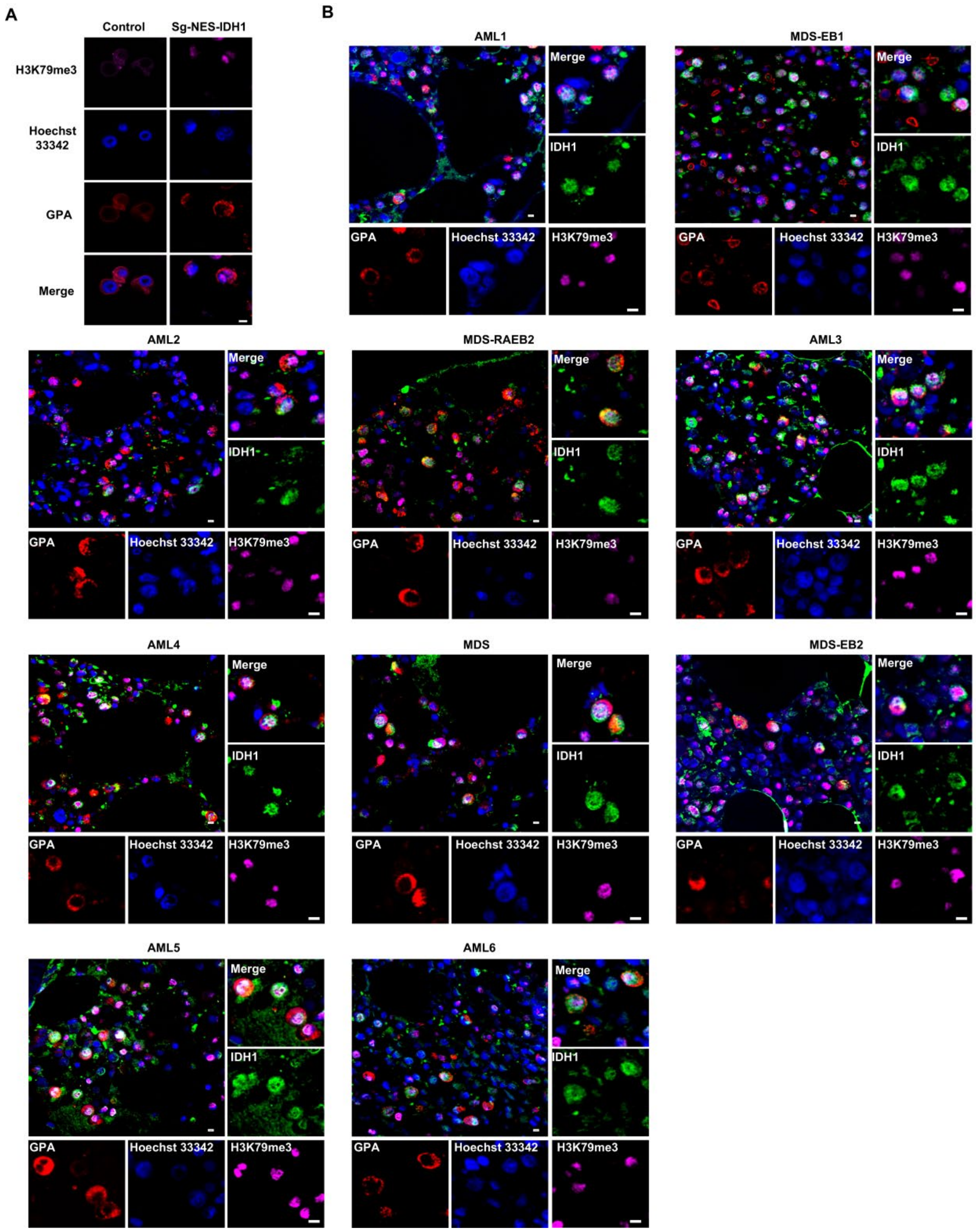


**Supplemental Figure 11.** The location of IDH1 and H3K79me3 in terminal erythroblasts of sg-NES-IDH1 HUDEP-2 cell line and AML/MDA patients.

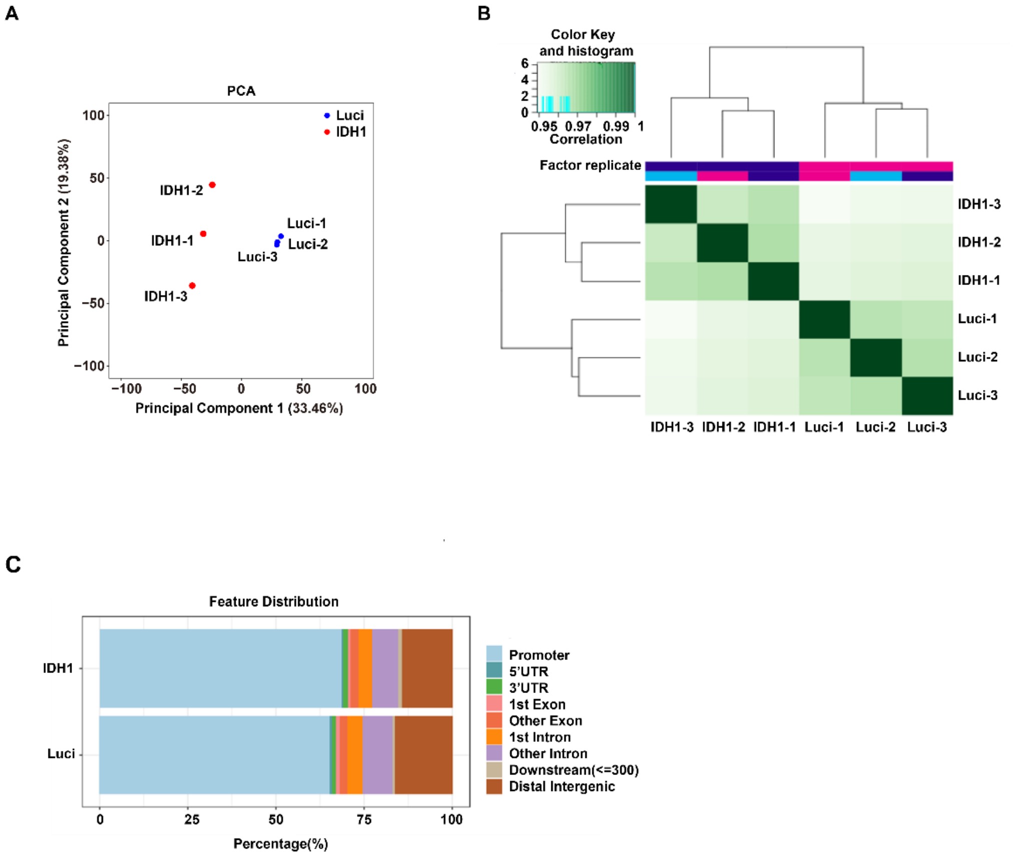


**Supplemental Figure 12.** ATAC-seq analysis.

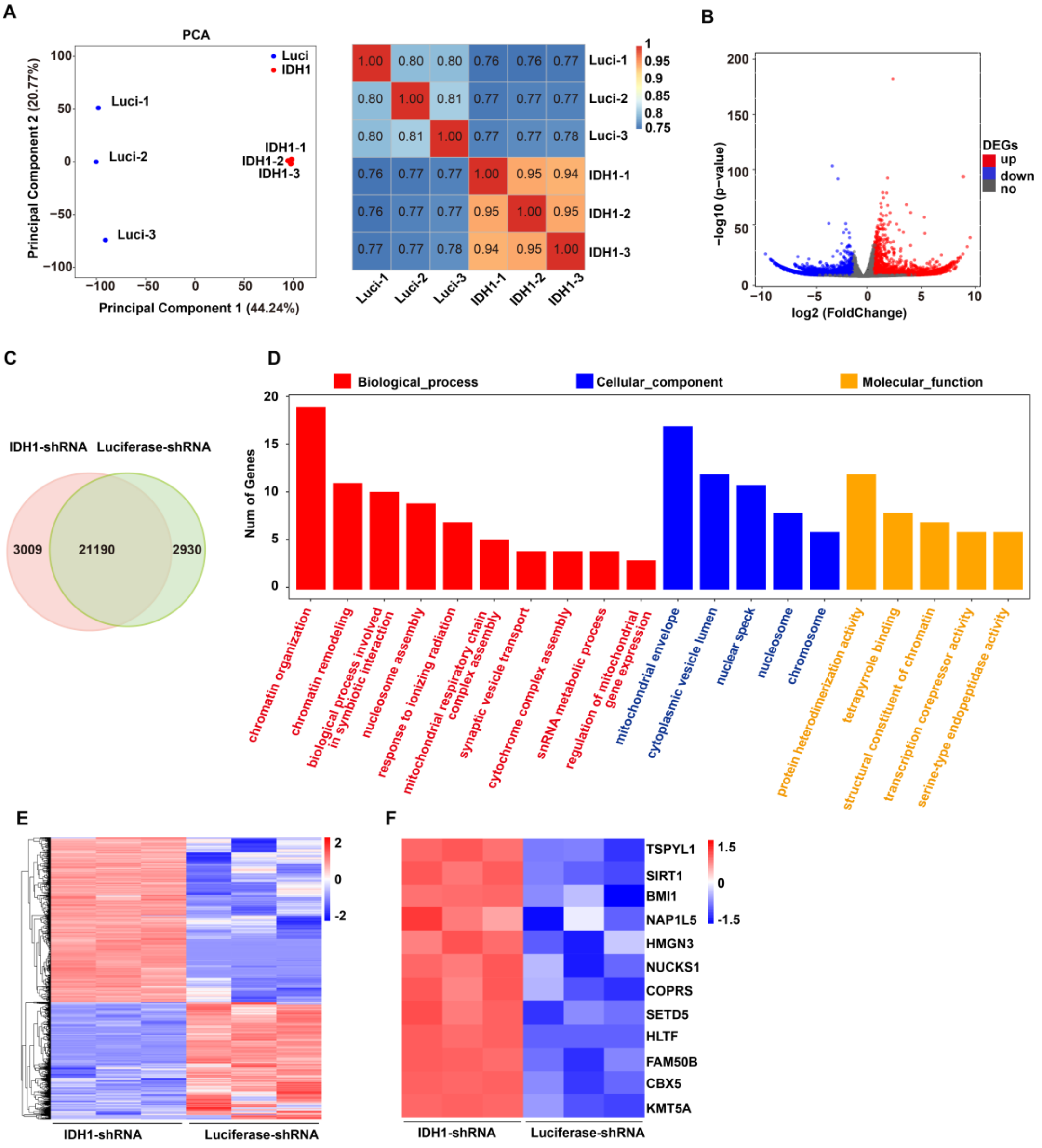


**Supplemental Figure 13.** RNA-seq analysis.

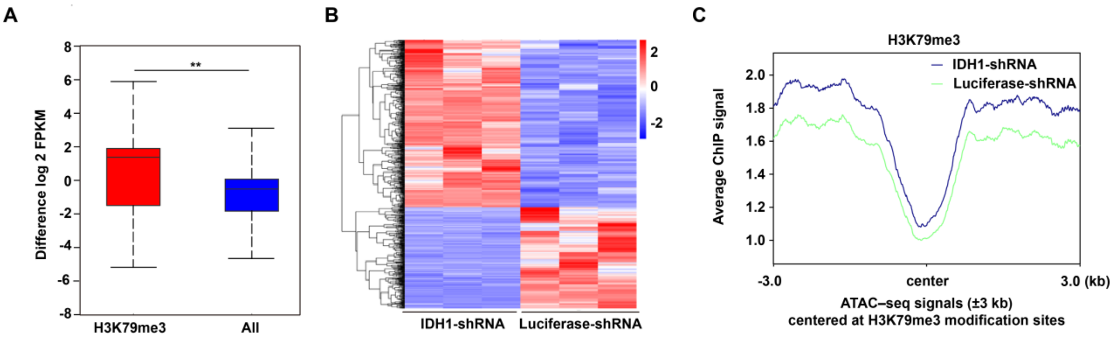


**Supplemental Figure 14.** Integrated analysis of ChIP-seq, ATAC-seq and RNA-seq

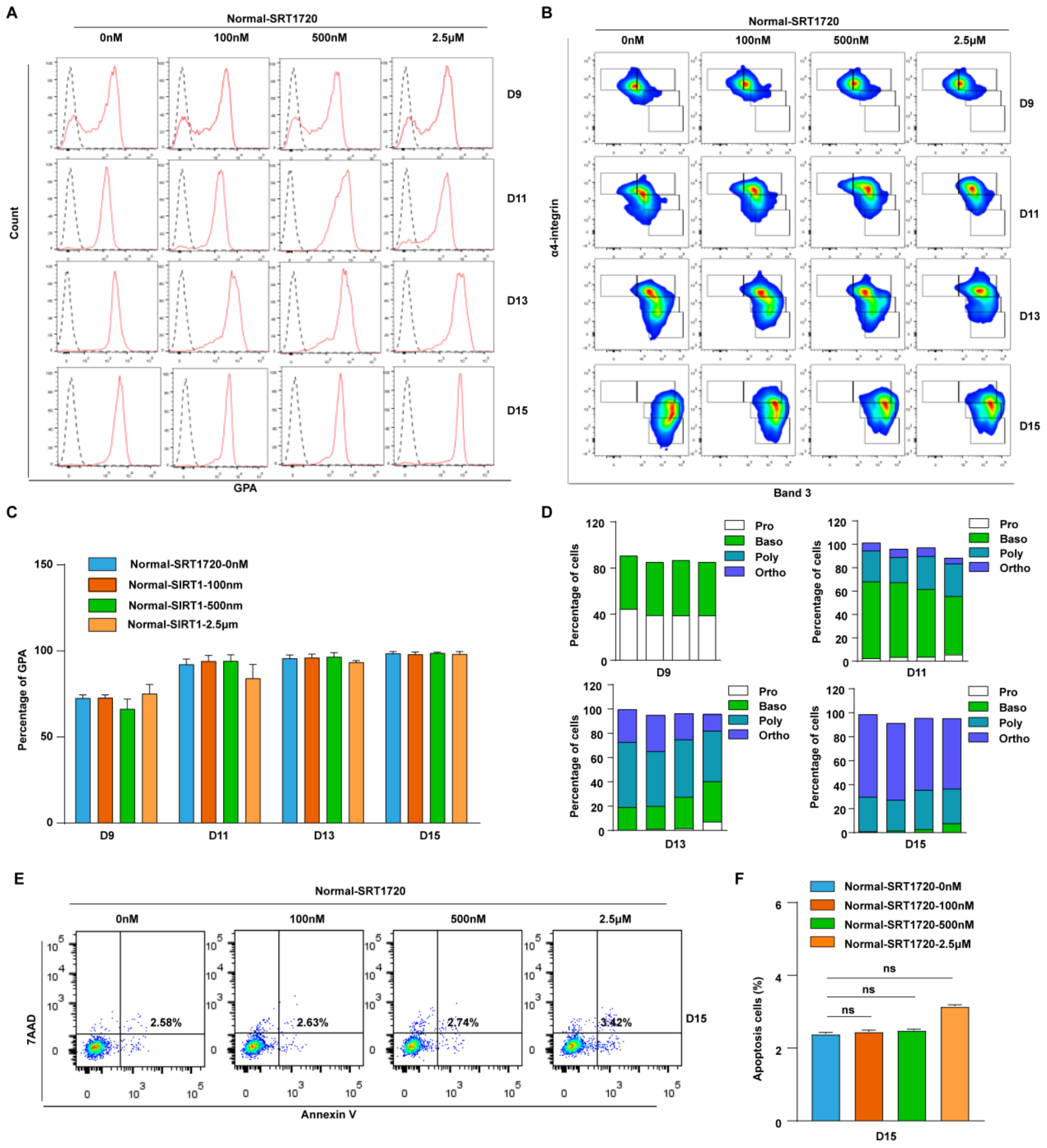


**Supplemental Figure 15.** Treatment with SIRT1 activator have no effet on cell differentiation and apoptosis of terminal erythroblasts.

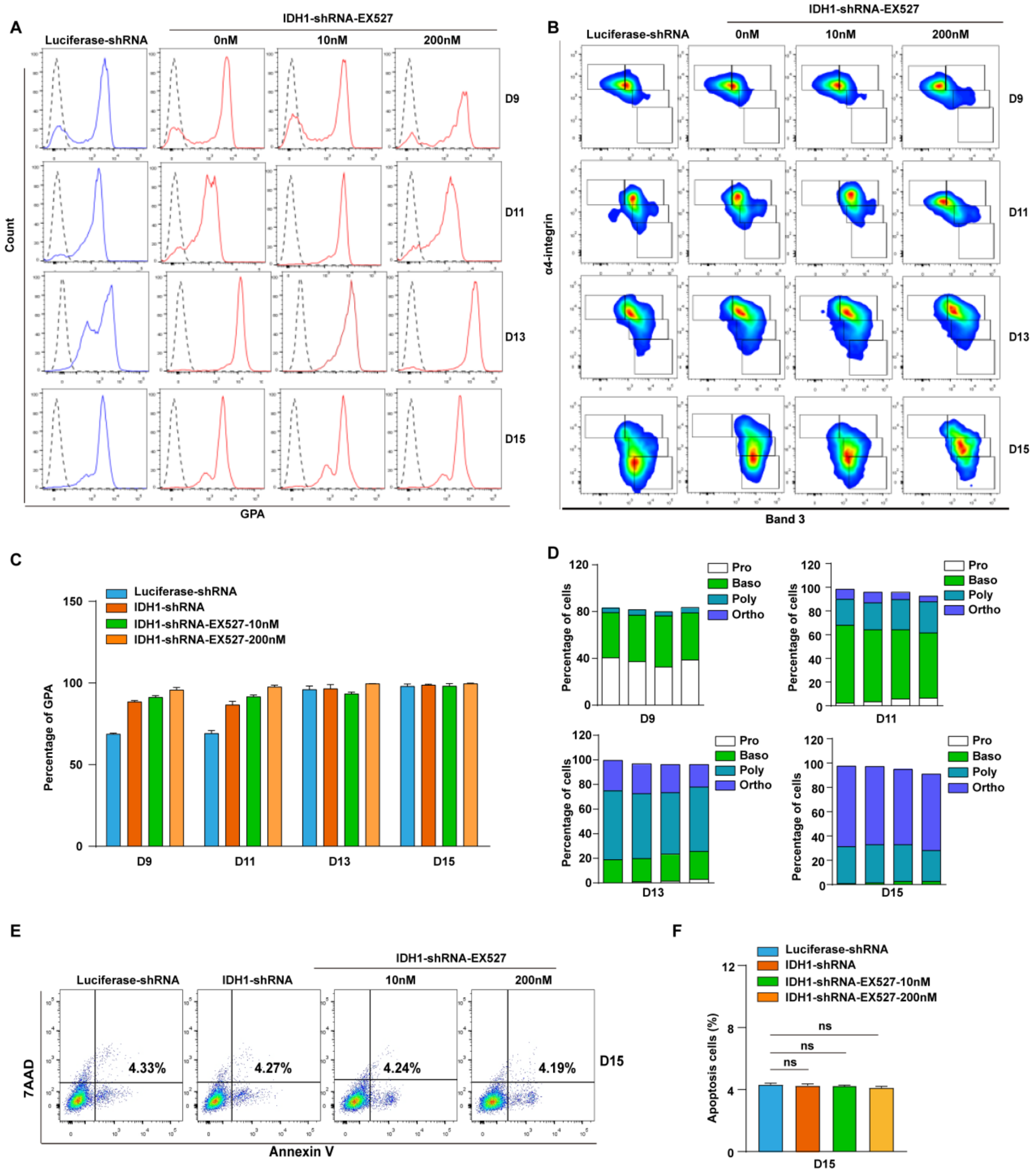


**Supplemental Figure 16.** Treatment with SIRT inhibitor have no effet on cell differentiation and apoptosis of terminal erythroblasts.

| No. | Sex | Age | Disease Type | Mutant Site | Frequency (%) |
|---|---|---|---|---|---|
| 1 | woman | 71 | AML | IDH1:NM_005896:exon4:c.C394T:p.R132C rs121913499 | 45.9 |
| 2 | woman | 16 | AML | IDH1:NM_005896:exon4:c.C394T:p.R132C rs121913499 | 45.81 |
| 3 | woman | 64 | MDS-EB1 | IDH1:NM_005896:exon4:c.C394T:p.R132C rs121913499 | 29.86 |
| 4 | woman | 48 | AML | IDH1:NM_005896:exon4:c.C394G:p.R132G rs121913499 | 26.98 |
| 5 | woman | 55 | AML | IDH1:NM_005896:exon4:c.C394T:p.R132C rs121913499 | 25.47 |
| 6 | woman | 71 | MDS -RAEB2 | IDH1:NM_005896:exon4:c.C394T:p.R132C rs121913499 | 18.72 |
| 7 | man | 56 | AML | IDH1:NM_005896:exon4:c.C394T:p.R132C rs121913499 | 13.16 |
| 8 | woman | 75 | MDS | IDH1:NM_005896:exon4:c.C394T:p.R132C rs121913499 | 12.97 |
| 9 | man | 46 | MDS -EB2 | IDH1:NM_005896:exon4:c.G395A:p.R132H rs121913500 | 5.71 |
| 10 | man | 59 | AML | IDH1:NM_005896:exon4:c.G395A:p.R132H rs121913500 | 1.93 |

**Supplemental Table 1.** Mutation analysis of MDS and AML patients with IDH1-mut.

| Primer | Sequence |
|---|---|
| IDH1 -shRNA1 | Forward:5'- CCGGCCTATCATCATAGGTCGTCATCTCGAGATGACGACCTATGATGATAGGTTTTTG-3' |
| | Reverse:5' - AATTCAAAAACCTATCATCATAGGTCGTCATCTCGAGATGACGACCTATGATGATAGG-3' |
| IDH1 - shRNA2 | Forward:5'- CCGGCCTTTGTATCTGAGCACCAAACTCGAGTTTGGTGCTCAGATACAAAGGTTTTTG-3' |
| | Reverse:5'- AATTCAAAAACCTTTGTATCTGAGCACCAACTCGAGTTTGGTGCTCAGATACAAAGG-3' |
| IDH1 - siRNA1 | GGCCCAAGCUAUGAAAUCATT. |
| IDH1 - siRNA2 | CCUGGUACAUAACUUUGAATT. |

**Supplemental Table 2.** The sequence for IDH1- shRNA and IDH1-siRNA.

| Primer | Sequence |
|---|---|
| IDH1 | Forward:5' - TGTGGTAGAGATGCAAGGAGA-3' |
| | Reverse:5' - TTGGTGACTTGGTTGGTG-3' |
| GAPDH | Forward:5'- CATGAGAAGTATGACAACAGCCT-3' |
| | Reverse:5'- AGTCCTTCCACGATACCAAAGT-3' |

**Supplemental Table 3.** The qRT-PCR primer sequence for IDH1 and GAPDH.
